# Comparing the Fate of Brain Metastatic Breast Cancer Cells in Different Immune Compromised Mice with Cellular Magnetic Resonance Imaging

**DOI:** 10.1101/2020.02.18.954693

**Authors:** Natasha N. Knier, Amanda M. Hamilton, Paula J. Foster

## Abstract

Metastasis is the leading cause of mortality in breast cancer patients, with brain metastases becoming increasingly prevalent. Studying this disease is challenging due to the limited experimental models and methods available. Here, we used iron-based cellular MRI to track the fate of a mammary carcinoma cell line (MDA-MB-231-BR) *in vivo* to characterize the growth of brain metastases in the nude and severely immune-compromised NOD/SCID/ILIIrg−/− (NSG) mouse.

Nude and NSG mice received injections of iron-labeled MDA-MB-231-BR cells. Images were acquired with a 3T MR system and assessed for signal voids and metastases. The percentage of signal voids and the number and volume of metastases were quantified. *Ex vivo* imaging of the liver, histology, and immunofluorescence labeling was performed.

On day 0, iron-labeled cells were visualized as signal voids throughout the brain. The percentage of voids decreased significantly between day 0 and endpoint. At endpoint, there was no difference in the number of brain metastases or tumour burden in NSG mice compared to nudes. Tumour volumes in nude mice were significantly larger than in NSG mice. Body images indicated that the NSG mice had metastases in the liver, lungs, and lymph nodes.

Characterization of the NSG and nude mouse is necessary to study breast cancer brain metastasis *in vivo*. Here, we show that the 231BR cell line grew differently in NSG mice compared to nude mice. This work demonstrates the role that imaging can play toward credentialing these models that cannot be done with other *in vitro* or histopathologic methods alone.

## Background

Breast cancer is a leading cause of death in women mainly due to the propensity of breast tumors to metastasize to regional and distant sites, such as the lymph node, lung, liver, bone and brain [1]. The incidence of brain metastases is increasing due to the introduction of more sensitive diagnostic methods and improved systemic therapies leading to improvements in extra-cranial control and survival [2–4]. Breast cancer is a disease with a number of subtypes and patients with metastatic ‘triple-negative’ breast cancer tend to develop brain metastasis at a high rate [5, 6]. For the HER2 amplified subtype, the frequency of brain metastasis has been reported to be as high as 50% [7].

Once a metastatic cancer cell arrives in the brain one of three things can happen: (1) it may die, (2) it may proliferate to form micrometastases, or (3) it may remain viable but dormant (‘non-proliferative’) [8, 9] If the solitary cells proliferate to form micrometastases, they may again experience one of three fates: (1) they may die, (2) they may continue to proliferate and form macrometastases, or (3) they may persist as “dormant” micrometastases, where dormancy is defined as a balance between proliferation and apoptosis within the cell population such that there is no net growth [9, 10]. The factors that tip the balance between dormancy and proliferation are poorly understood. Both dormant single cells and dormant micrometastases are believed to be sources of cells that contribute to tumor recurrence [11]. Dormant cancer cells also present a substantial therapeutic problem; since they are quiescent, they are non-responsive to current therapies which target proliferating cells [8].

Currently, only a handful of models specific to breast cancer brain metastasis have been described and even fewer allow for *in vivo* investigation of cancer cell dormancy. Both murine and human cancer cell lines have been developed to mimic as many steps in the metastatic cascade as possible. A well characterized murine breast cancer cell line is the 4T1-BR5 cell line, a highly tumorigenic and invasive cell line that has undergone multiple rounds of selection to preferentially grow in the mouse brain [12]. There are several advantages to using a murine breast cancer cell line to study brain metastasis, as growth and maintenance is easy and relatively inexpensive, and it can be grown in immune-competent mice, which is of particular interest for studying this disease in a way that recapitulates the tumor microenvironment [13].

A small number of human breast cancer brain metastatic cell lines have also been developed, including the commonly used MDA-MB-231BR (231BR), MDA-MB-231BR-HER2, MA11, JIMT1-BR3, SUM190-BR3 cell lines. The MDA-MB-231BR cell line has been particularly well characterized for studying the progression of brain metastases in nude mice. Nude mice are the most commonly used immune-deficient strain for models which use human cell lines. They have a genetic mutation that causes a deteriorated or absent thymus, resulting in a lack of T cells [14]. The 231BR cell line grows selectively in the brain of nude mice, without metastatic growth seen in other distant organs [15]. Human breast cancer cell lines are one of the most widely used models to study the metastatic growth of cancer *in vitro* and *in vivo*, as they have been used to provide extensive insight into the characteristics of human cells and can be used for high throughput screening of various drugs [16]. There are, however, significant limitations to using human and murine breast cancer cell lines, as the quick progression *in vivo* can limit opportunities for adequate therapeutic testing, and the growth of these cells *in vitro* prior to establishment in a mouse can cause changes in the genetic composition due to clonal selection [17]. Many groups have shown that these genetic changes result in these models failing to maintain tumor heterogeneity, which is now recognized as a critical element for developing personalized treatments [18–20]. Studying breast cancer with immunocompetent mice also has limitations, as tumor latency and growth can be variable and slow [21]. There is also increased difficulty in establishing human tumors successfully in these mice. Immune-compromised mice lack a comparable tumor microenvironment to clinical tumors and fail to provide a realistic result of interactions with the natural immune response, particularly in the case of studying anticancer therapeutics [22].

In more recent years, researchers have moved towards studying breast cancer and its subtypes with patient-derived xenografts (PDX), which allow for the growth of human primary breast cancer tissue that has been recently resected from consenting patients into immune-compromised rodents [23]. NOD/SCID/ILIIrg^−/−^ (NSG) mice are the preferred strain of mice to engraft a PDX, as they are highly permissive to growing breast cancer metastasis, and resembles the metastatic pattern seen in human patients. NSG mice are the most immunodeficient mouse strains to date, lacking T cells, B cells, NK cells, and have defective macrophages and dendritic cells [24–26]. Even fewer PDX models have been developed specific to breast cancer brain metastasis, including the novel F2-7 and E22-1 PDX cell lines [27], as well as others who have established low passage PDX models of breast cancer brain metastasis and those that implant fresh tumor tissue directly into the rodent brain [28].

The D2.0R and related D2A1 mouse mammary tumor cells have been well characterized as model systems for tumor cell dormancy. In mice, D2.0R/R cells invade distant metastatic sites, remain as single quiescent cells for prolonged periods of time, and occasionally proliferate into metastatic tumors. D2A1/R cells, have a much shorter dormancy period before forming metastases in the lung, liver, and other organs [29]. A human breast cancer cell line, MCF-7 has also been studied to examine the mechanisms of dormancy, however it is poorly metastatic [30].

For all of these brain metastasis models – cancer cell growth, metastasis and dormancy are typically studied using methods such as histology, flow cytometry, immunohistochemistry, and fluorescent microscopy [31]. While these techniques provide useful information about molecular and cellular markers and morphology, they are limited to studying this disease after endpoint has been reached. There is a clear need to characterize brain metastatic models of breast cancer *in vivo*, which can be accomplished through use of imaging modalities and cell tracking techniques.

Cellular magnetic resonance imaging (MRI) combines the ability to obtain high resolution MRI data with the use of iron oxide nanoparticles for labeling specific cells, thereby enhancing their detectability [32, 33]. The presence of the iron in cells causes a distortion in the magnetic field and leads to abnormal signal hypo-intensities in iron-sensitive images (T2*-weighted images are most often used). Areas containing iron labeled cells therefore appear as regions of low signal intensity on MRI images, creating negative contrast [34]. We have previously shown that it is possible to use cellular MRI to track iron-labeled 231BR cancer cells in the nude mouse brain [35–37]. Proliferative cancer cells lose the iron label over time; as the cells divide the particles are apportioned to daughter cells and eventually some cells contain too little iron to be detectable by MRI. As brain metastases form, changes to the tissue result in the tumor appearing brighter than the surrounding brain in MRI. Nonproliferative cancer cells retain the iron particles and can be detected over long periods of time [37].

In this study, we used these cellular MRI techniques to characterize and compare the growth of 231BR brain metastases and the persistence of iron-retaining cancer cells in nude and NSG mice. Our goal was to evaluate the NSG mouse as a model for breast cancer brain metastasis and dormancy.

## Methods

### Cell Culture and MPIO Labeling Procedure

Brain trophic human breast cancer cells (MDA-MB-231BR) expressing green fluorescent protein (GFP) were maintained with Dulbecco’s modified Eagle’s medium containing 10% fetal bovine serum (complete DMEM) at 37°C and 5% CO_2_. 2×10^6^ of these cells were seeded and allowed to adhere for 24 hours. To iron label these cells, cells were supplemented with 25 μg Fe/mL of micron-sized iron oxide particles (MPIO) (0.9 μm diameter, ~63% magnetite, labeled with Flash Red; Bangs Laboratory, Fishers, IN, USA) in 10mL of complete DMEM in a T75-cm^2^ flask. Following this, cells were washed once with phosphate buffered solution (PBS) within the flask and trypsinized with 0.25% Trypsin-EDTA. After, cells were harvested and washed three additional times with PBS in the flask to thoroughly remove unincorporated MPIO prior to cell injections. Cell viability was assessed and calculated using the Trypan blue exclusion assay. To visualize MPIO labelling, labeled cells were affixed to a glass slide with a ThermoFisher Cytospin 4 cytocentrifuge and fixed with a Methanol/Acetic acid solution. Slides were then stained with a Perl’s Prussian Blue (PPB) solution and counterstained with Nuclear Fast Red. Slides were dehydrated with increasing concentrations of ethanol, cleared with xylene, and coverslipped with a xylene-based mounting medium. These PPB-stained slides were examined to assess the localization of MPIO within the cell and to determine the labeling efficiency using a Zeiss AXIO Imager A1 Microscope (Zeiss Canada, Toronto, ON, Canada). Iron oxide nanoparticles appear dark blue, and cells appear light pink in colour.

### Animal Model and Work Flow

All animals were cared for in accordance with the standards of the Canadian Council on Animal Care, under an approved protocol of the University of Western Ontario’s Council on Animal Care and housed in a pathogen-free barrier facility. Female nude mice (nu/nu Foxn1, aged 6-8 weeks, from Charles River Laboratories, Wilmington, MA) and female NSG mice (NOD.*Cg-Prkdc*^*scid*^*Il2rg*^*tm1Wjl*^/*SzJ*, 6-8 weeks, Humanized Mouse and Xenotransplantation Facility, Robarts Research Institute, University of Western Ontario, London, ON) were first anesthetized with isoflurane (2% in 100% oxygen). Nude (n=10) and NSG mice (n=10) were then injected with a suspension of 1.5×10^5^ MPIO-labeled MDA-MB-231BR/GFP+ cells in 100μL of sterile saline and 15% Vevo MicroMarker microbubble solution (FUJIFILM, VisualSonics Inc., Toronto, ON, Canada). Cell suspension was loaded into a 100μL Hamilton syringe with a 30G needle. Cells were administered by slow intracardiac injection into the beating left ventricle of the heart with ultrasound imaging guidance on a Vevo 2100 ultrasound system (FUJIFILM, VisualSonics Inc., Toronto, ON, Canada). Cancer cell arrest was evaluated with MRI on the day of the injection (Day 0) and used to determine whether mice had a successful intracardiac injection. Absence of signal voids in the brain indicated an unsuccessful injection, resulting in exclusion from the remainder of the study. Nude mice with successful injections had MRI performed at day 21 and day 32 post-injection. NSG mice with successful injections had MRI performed at day 21 post-injection (Figure 1).

**Figure. 1.**
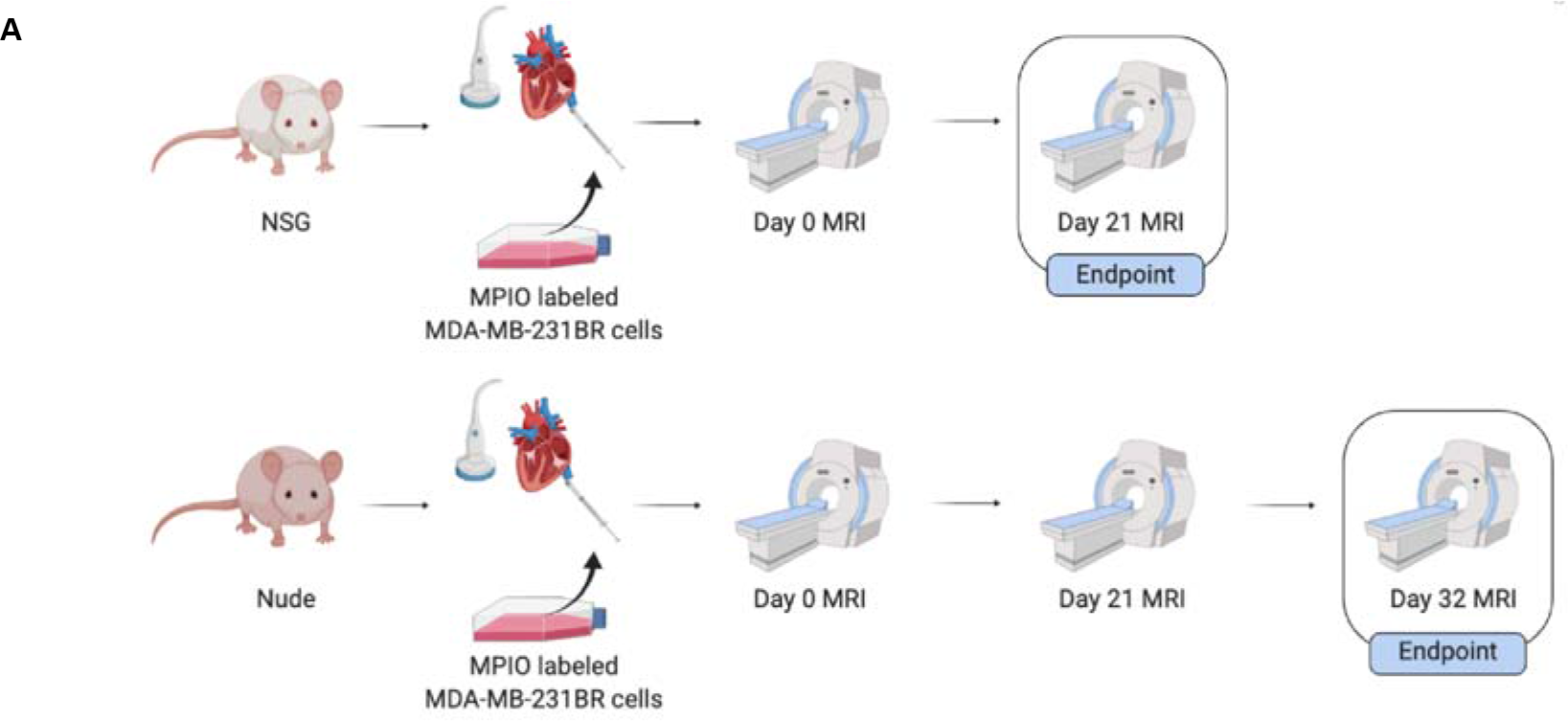
Experimental design for the NSG and nude mouse model.

Additionally, 1 nude mouse and 3 NSG mice received intracardiac injections with 1.5×10^5^ MPIO labeled MD-MB-231BR/GFP+ cells and intraperitoneal injections of 200μL of gadopentetate dimeglumine (Magnevist, Schering, US; 0.5 mmol/mL) on Day 0 to assess cancer cell arrest in the liver. These mice were sacrificed on Day 0 by isoflurane overdose approximately 40 minutes after intraperitoneal administration. These mice were then exsanguinated, and brains and livers were removed for *ex vivo* imaging.

### Magnetic Resonance Imaging

All brain and body MRI examinations were acquired on a 3.0 Tesla GE MR750 clinical MR scanner (General Electric, Mississauga, ON, Canada) using a custom-built gradient coil (inner diameter = 17.5 cm, gradient strength = 500 mT/m, and peak slew rate = 3000 T/m/s]). Brain and ex vivo liver images used a custom-built solenoidal mouse brain radiofrequency (RF) coil. Body images were acquired using a 4.3 × 4.3 cm dual tuned ^1^H/^19^F surface coil (Clinical MR Solutions, WI, USA), originally built for imaging small ROIs in humans. Mice were anesthetized with 2% isoflurane in 100% oxygen administered through a nose cone. *In vivo* brain and body images were acquired using a 3D balanced steady-state free precession (bSSFP) sequence [Fast Imaging Employing Steady State Acquisition (FIESTA) on a GE system] that has been optimized for simultaneous detection of signal voids produced by iron-loaded cells and hyperintense metastases. This permitted the assessment of both cell arrest and retention as well as the number and volume of metastases throughout the mouse brain. Brain images for nude mice were acquired on day 0, day 21, and day 32. Brain imaging for NSG mice occurred on day 0 and day 21. Scanning parameters were as follows: resolution = 100 × 100 × 200 μm, repetition time (TR) = 7 ms, echo time (TE) = 3.5 ms, bandwidth (BW) = 20.83kHz, flip angle (FA) = 35 degrees, signal averages = 2, phase cycles = 8, matrix = 150 × 150, scan time = approximately 33 minutes per mouse. Body images were acquired on Day 20. Body imaging parameters were as follows: resolution = 200 × 200 × 200 μm, repetition time (TR) = 4.7 ms, echo time (TE) = 2.3, bandwidth (BW) = +/−31.25 kHZ, flip angle (FA) = 35 degrees, signal averages = 2, phase cycles = 8, matrix = 250 × 250, scan time = approximately 20 minutes. *Ex vivo* liver imaging parameters were acquired using a spoiled gradient echo sequence that also allows for the detection of signal voids throughout the liver. *Ex vivo* liver samples were immersed in tubes containing Fluorinert™ FC-40 Electronic Liquid (3M, St. Paul’s, MN, USA), a fluorocarbon liquid which produces a black background in proton images. Images were taken of ex vivo livers acquired on day 0 for both strains, day 21 for the NSG mouse, and day 32 for the nude mouse. The scanning parameters for ex vivo liver images were as follows: resolution = 100 × 100 × 200 um, repetition time (TR) = 43 ms, echo time (TE) 4.844 ms, bandwidth (BW) = 31.25kHz, flip angle (FA) = 60 degrees, signal averages = 1, phase cycles = 8, matrix = 250 × 250, scan time = approximately 23 minutes.

### Image Analysis

Images were analyzed using open-source OsiriX image software (Pixmeo, SARL, Bernex, Switzerland), version
10.0.4 and Horos image software, version 3.3.5. Brain images were evaluated for successful cell delivery by assessment of signal voids on day 0. To quantify cancer cell arrest throughout the brain, the number of black pixels within the total brain volume was determined from day 0 MRI images. The brain was manually segmented as a region of interest and then a threshold value was determined based on the mean pixel intensity value of a representative signal void within the brain region ± 2 standard deviations. The total number of black pixels under this threshold value within the entire brain volume signal was obtained from a pixel intensity histogram. To quantify brain metastases at day 21 and day 32, tumors were counted in all image slices by a single observer. Each tumor was manually segmented, and the tumor volumes were reconstructed using the OsiriX and Horos volume algorithm. To calculate brain tumor burden at day 21 and day 32, an ROI was drawn around the outline of the brain in each slice, and a 3D reconstruction using the OsiriX and Horos volume algorithm provided a quantification of the total brain volume. The total volume of all segmented tumors was then determined by adding each individual tumor volume measurement and calculating the percentage of the total brain volume occupied by tumors. Body images were qualitatively assessed for the presence of metastases in the liver, lung, and lymph nodes. All quantitative values were presented as the mean ± standard error. Statistical analysis was performed using Welch’s *t* tests on GraphPad Prism version 8 software (GraphPad, San Diego, CA).

### Histology and Immunohistochemistry

At each strain’s respective endpoints, mice were euthanized by isofluorane overdose and perfused with 4% paraformaldehyde. Brains were removed and placed into paraformaldehyde for another 24 hours. Fixed brains were processed, paraffin embedded and then cut into 6 or 8 um sections. Sliced sections were deparaffinized and stained with either hematoxylin and eosin (H&E) or immunofluorescently labelled for Ki67.

### Hematoxylin and eosin (H&E) staining

Sections were washed briefly in distilled water, stained in Harris hematoxylin solution for 5 minutes and differentiated in 1% acid alcohol for 30 seconds. After washing in 0.2% ammonia for 5 minutes, sections were counterstained in eosin-phloxine solution for 30 seconds; dehydrated through 95% alcohol, 2 changes of absolute alcohol, and 5 minutes each. Sections were then dehydrated and cleared through 95% ethyl alcohol, absolute alcohol and xylene, and mounted with resinous medium.

### Immunofluorescent labelling

Ki-67 immunostaining was performed using a rat anti-Ki-67 antibody (1:400 dilution; Catalog #14-5698-82, Invitrogen). All sections were permeabilized with 0.2% Triton X-100 in PBS for 15 minutes and non-specific protein binding was then blocked by incubation in a commercial blocking agent (ab156024, Abcam) for 1 hour at room temperature. Sections were then incubated with the Ki-67 primary antibody in commercial antibody dilutant (ab64211, Abcam) at room temperature for 1 hour. Negative controls (without addition of primary antibody) were performed on adjacent sections. Unbound primary antibody was washed away through three 5-minute exchanges of 1×PBS. Sections were then incubated with an anti-rat Alexa Fluor-488 secondary antibody (1:300 dilution; Catalog #A-11006, Invitrogen) for 1 hour. Unbound secondary antibody was washed away through three 5-minute exchanges of 1×PBS. Finally, nuclei were counterstained with Hoechst 33258 for 5 minutes and rinsed sections cover slipped for microscopic examination.

## Results

### *In Vitro* Studies

#### Cell Labelling

MDA-MB-231BR cells were efficiently labeled with MPIO, as demonstrated by the Perl’s Prussian blue staining of cells shown in Figure 2a; cancer cells appear pink, and the intracellular iron is blue. Labelling efficiency of 96.33±1.20% was achieved with 97.33±0.88% cell viability.

**Figure. 2.**
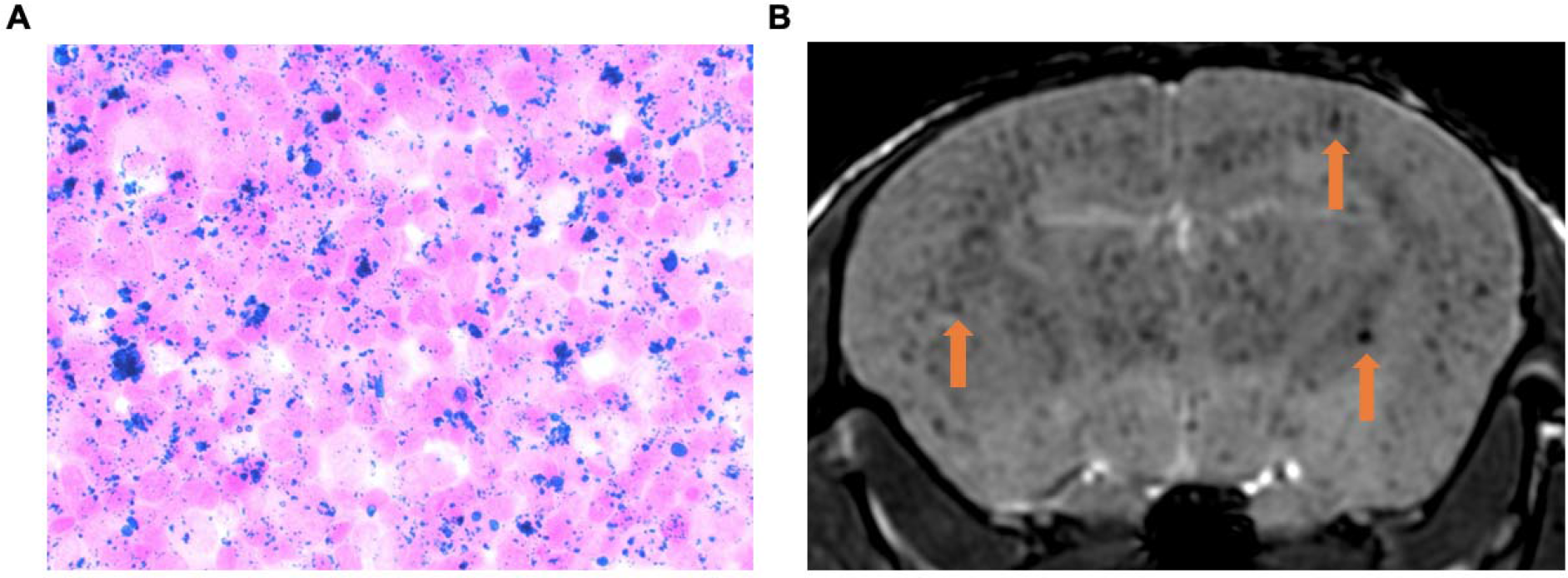
Iron labeling and injection of 231BR cells. (A) Perl’s Prussian Blue stain showing intracellular iron (blue, PPB) detected within MDA-MB-231BR cells (pink, Nuclear fast red) *in vitro*. (B) Representative day 0 image showing visualization of successful iron-labeled cancer cell delivery by intracardiac injection as regions of signal void (orange arrows)

### *In Vivo* Studies

#### Experimental Endpoint

In our experience, intracardiac injection of 1.5×10^5^ 231BR cells in nude mice results in brain tumor burden, significant weight loss and neurological impairments which leads to a requirement for euthanasia at 28-34 days post injection. In this study the nude mice reached experimental endpoint on day 32. In contrast, NSG mice could only be studied until day 21, at which time they had reached a weight loss greater than 15% of their body weight, resulting in extreme cachexia and anorexia. Additionally, these mice showed severe signs of neurological impairment, resulting in the inability to perform normal functions and exhibiting paralysis and circling. Post-mortem examination also revealed significant liver tumor burden, which also contributed to the early, unexpected endpoint.

#### Imaging Cell Arrest and Retention

On day 0, bSSFP brain images confirmed the successful intracardiac injection of MPIO-labeled cancer cells in all mice; iron-labeled cancer cells appeared as distinct regions of signal void throughout the brain due to their initial arrest in brain vasculature (Figure 2b). Figure 3a shows representative day 0 images for each mouse strain. When quantifying the percentage of the brain consisting of black signal voids, approximately 4.69%±1.71 and 6.26%±1.86 of the brains of NSG and nude mice contained arrested cancer cells, respectively. At each strain’s endpoint, the number of black pixels was again determined, with percentages significantly reduced to approximately 1.89%±0.57 in NSG mice (day 21) and 2.45%±0.48 in nude mice (day 32). While the number of arrested cancer cells significantly differs from day 0 to endpoint for both NSG (p = 0.03) and nude mice (p = 0.02), there was no significant differences between strains at each time point (Figure 3b).

**Figure. 3.**
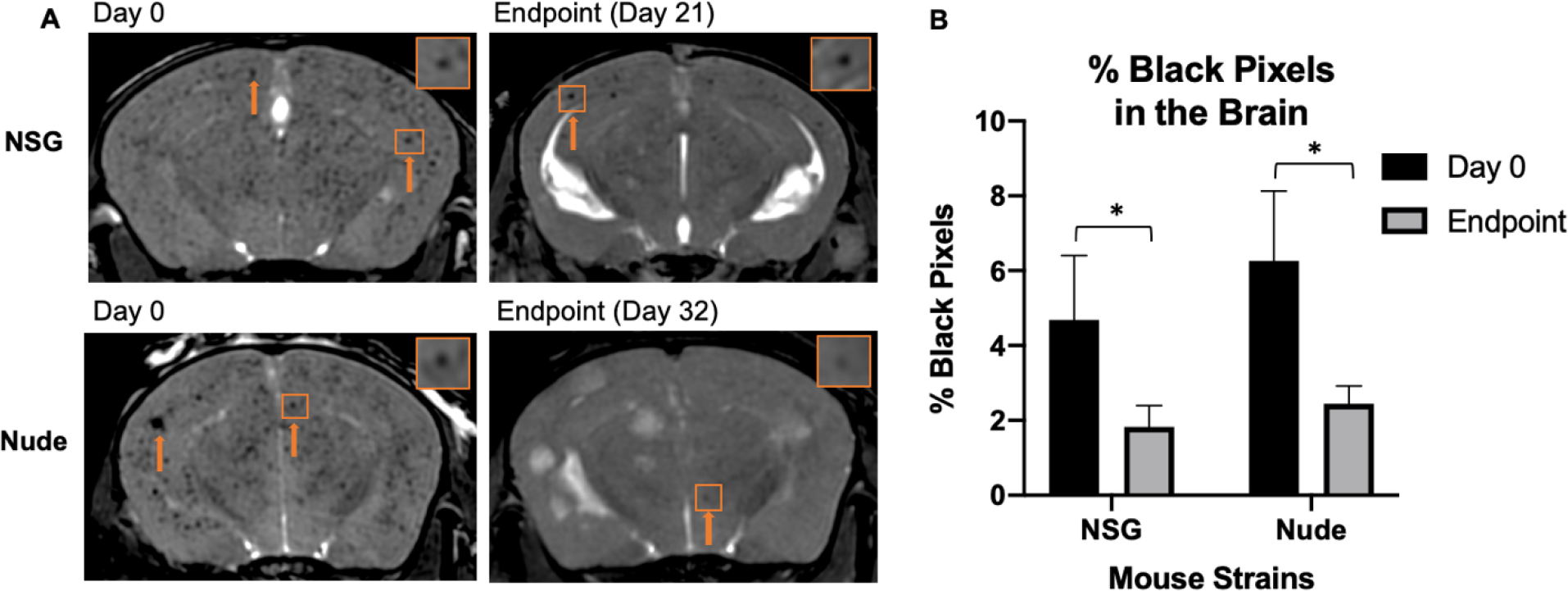
Cancer cell arrest and clearance. (A) Balanced steady-state free precession images showing initial cancer cell arrest at day 0 and clearance and retention at endpoint. The orange arrows indicate regions of signal void. (B) The percentage of black pixels, representing regions of signal void was not significantly different between strains, however there was a significant decrease within strains from day 0 to endpoint, suggesting similar cancer cell arrest and clearance between nude and NOD/SCID/ILIIrg−/− mice

#### Day 21 MRI

On day 21, metastases were detectable in bSSFP brain images as regions of signal hyperintensity in both strains of mice (Figure 4a). On day 21, the mean number of MRI-detectable metastases was significantly higher in NSG mice (63.70±5.37) compared to nude mice (15.33±4.29) (Figure 4b). While the nude mice had fewer tumors, they were significantly larger. The mean volume of tumors in nude mice was 0.06 mm^3^±0.01, compared to 0.04 mm^3^±0.003 for NSG mice (Figure 4c). This mean volume was based on a total number of 637 tumors in all NSG mice and 72 tumors counted in all nude mice. At day 21, the mean tumor burden was significantly higher for NSG mice (2.39 mm^3^±0.18) compared to nude mice (0.82 mm^3^±0.35) (Figure 4D).

**Figure. 4.**
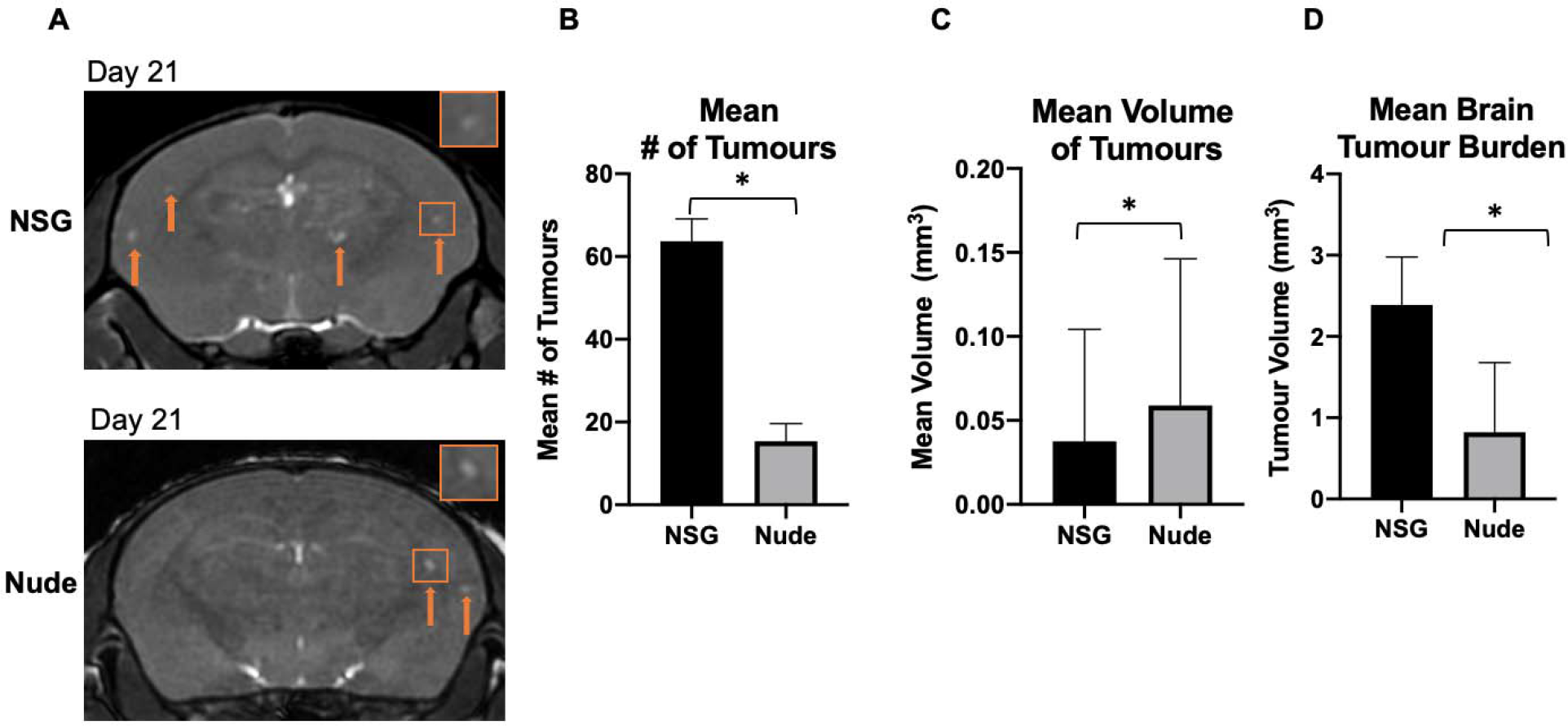
Day 21 visualization and quantification of brain metastases. (A) Balanced steady-state free precession images showing brain metastases in both strains of mice as regions of signal hyperintensity (orange arrows) at day 21. (B) There were significantly more brain metastases on average detected with MRI in NSG mice than nude mice at day 21. (C) The mean volume of tumors present in the brain was significantly greater in nude mice than NSGs. (D) The brain tumor burden of NOD/SCID/ILIIrg−/− mice was significantly higher than in nude mice at day 21. Data is presented as mean +/− SEM. *indicates p < 0.05

#### Day 32 MRI

Nude mice were followed out their endpoint of day 32 and imaging analysis was compared to the endpoint data for the NSG mice (Figure 5). The mean number and volume of brain metastases in nude mice increased between days 21 and 32. On day 32 the mean number of brain metastases in nude mice was 39.60±10.99. When comparing the mean number of metastases for NSG and nude mice at their respective endpoints there was no longer a significant difference (p = 0.07) (Figure 5b).

**Figure. 5.**
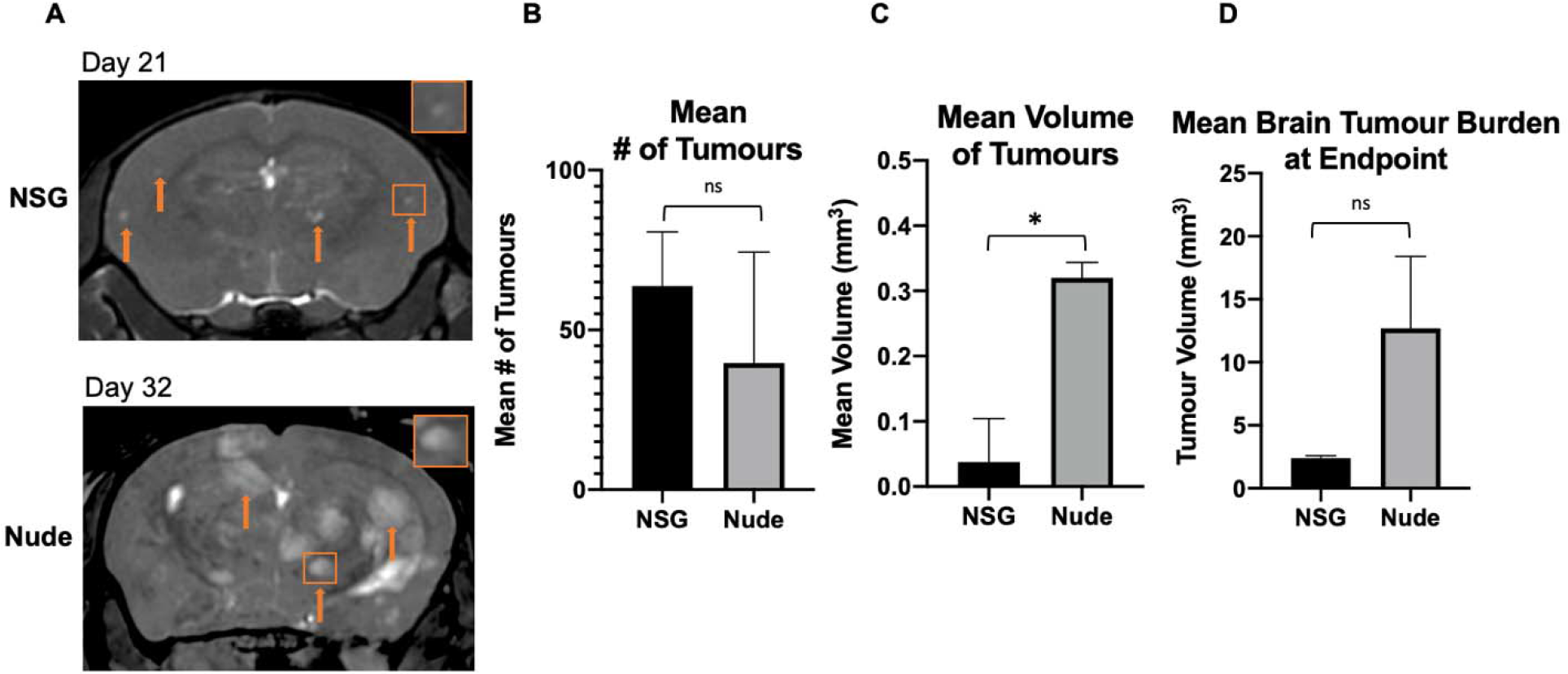
Endpoint visualization and quantification of brain metastases. (A) Magnetic resonance images of both NOD/SCID/ILIIrg−/− and nude mice at each strain’s respective endpoints (Day 21 for NOD/SCID/ILIIrg−/−, Day 32 for nude) showing regions of signal hyperintensity where brain metastases have developed (orange arrows). (B) There was no significant difference between the number of brain metastases detected with magnetic resonance imaging between strains at endpoint. (C) The tumors in the nude mice were significantly larger in volume than those in the NOD/SCID/ILIIrg−/− mice. (D) The total tumor burden within the brain at endpoint in both nude and mice was not significantly different. Data is presented as mean +/− SEM. *indicates p < 0.05

On day 32 the mean volume of brain metastases in nude mice was 0.32±0.23 mm^3^, compared to 0.04±0.03 mm^3^ on day 21. This was significantly higher (p = <0.0001) than the mean volume of brain metastases at the NSG endpoint (Figure 5c). Accordingly, nude mice had significantly more tumor burden than NSG mice (Figure 4d), which was a reversal from the previous timepoint at day 21. Due to the earlier endpoint of NSG mice but lesser brain tumor burden, body MRI was acquired for an NSG mouse at day 20. Notably, there was significant tumor burden detected by MRI within the liver, lungs (Figure 6A), and lymph nodes (Figure 6b) of the NSG mice. Nude mice had no MRI-detectable tumors within the body outside the brain (Figure 6c).

**Figure. 6.**
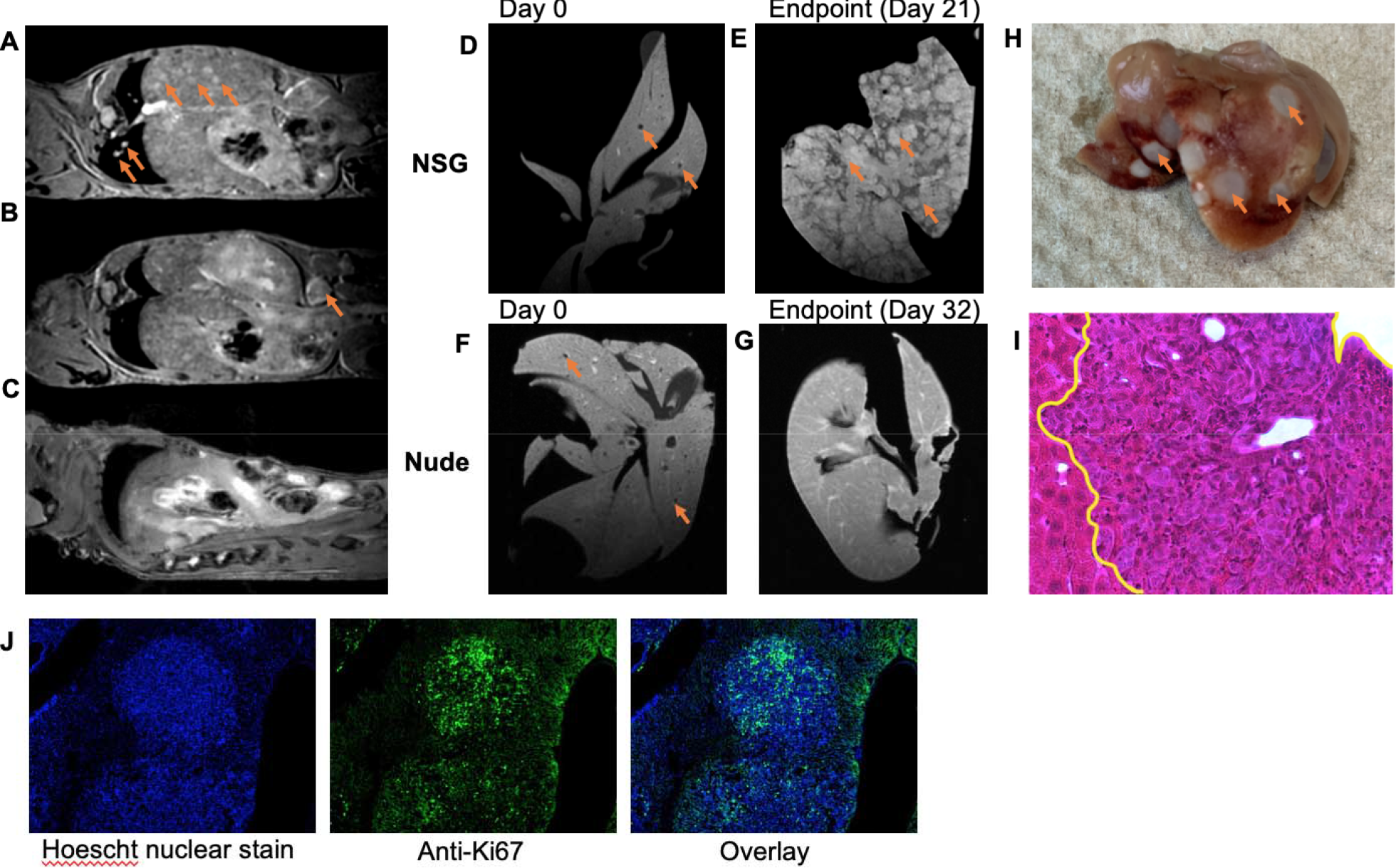
Body and liver imaging with correlative histology and immunofluorescence. (A) Body imaging of NOD/SCID/ILIIrg−/− mouse allowed for detection of significant tumor burden in the liver, lungs. (B) Tumors were detected in the lymph nodes of NOD/SCID/ILIIrg−/− mice with magnetic resonance imaging. (C) No tumors were detected with magnetic resonance imaging in the nude mouse body. (D) Spoiled gradient echo sequence image showing initial arrest of cancer cells in *ex vivo* liver at Day 0 for NOD/SCID/ILIIrg−/− mouse (E) Spoiled gradient echo sequence image of *ex vivo* liver showing regions of hyperintensity where liver metastases have developed in the NOD/SCID/ILIIrg−/− mouse. (F) Day 0 spoiled gradient echo sequence image of *ex vivo* liver of from a nude mouse showing signal voids, representing arrested cancer cells. (G) Spoiled gradient echo sequence image of Day 32 (endpoint) *ex vivo* liver of a nude mouse with no magnetic resonance detectable metastases (H) Photo of a representative *ex vivo* liver from an NOD/SCID/ILIIrg−/− mouse showing tumors on the exterior of the organ. (I) H&E-stained section showing liver metastases that had developed in the NOD/SCID/ILIIrg−/− mouse (outlined in yellow). (J) Ki67 staining showing a highly proliferative tumor within the NOD/SCID/ILIIrg−/− liver

#### Ex Vivo Liver MRI

Discrete signal voids were visible in the day 0 images of *ex vivo* livers from each strain of mouse, suggesting the arrest of iron labeled 231BR cells throughout the liver (Figure 6d,e). Ex vivo liver mages obtained from mice at endpoint showed numerous regions of abnormal high signal intensity associated with liver metastases, confirming the *in vivo* image results (Figure 6f). A photo of a representative liver sample clearly shows the liver metastases on the surface (Figure 6h). No regions of abnormal signal hyperintensity were observed in ex vivo images of the nude mouse liver (6g). H&E stained sections of the liver tissue from NSG mice confirmed the presence of metastases (6i) and ki67 staining showed that they were highly proliferative (6j).

## Discussion

This work demonstrates for the first time the use of *in vivo* longitudinal MRI based cell tracking to compare murine models of brain metastatic breast cancer. We show that there are significant differences in tumor progression for 231BR cells in NSG versus nude mice. While the initial arrest, clearance, and retention of iron-labeled cells was similar, brain metastases developed more quickly in NSG mice and NSG mice developed substantial body tumor burden, particularly in the liver, leading to an earlier endpoint.

The 231BR cell line was developed by Yoneda et al [15]. and is a brain-colonizing subline of the metastatic triple-negative MDA-MB-231 human breast cancer cell line, which was isolated by six repeated cycles of intra-cardiac injection and harvesting from brain metastases grown in nude mice. In our lab, the intra-cardiac injections are performed using ultrasound (US) guidance. Of note, was our observation by US of a thicker heart muscle surrounding the left ventricle of the NSG mouse. This anatomical observation made intracardiac injections more difficult, as the ventricular space appeared smaller with US. The observation of signal voids in brain images acquired on day 0 was used to identify successful delivery of cells to the brain. In this study, all mice imaged had successful injections and were included in the study. Delivery of cells to a specific organ is related to the cardiac output that is delivered to that organ. The cardiac output to the brain of a mouse is ~9.5% [38]. Here, we injected 150,000 cells into the left ventricle of the heart, and therefore we can expect ~14,250 cells to be successfully delivered to the brain. Only a percentage of these cells will arrest; previous studies, however, have shown that less than 1% of cells are retained in the microcirculation of the brain after 2 hours post-injection [38]. In previous studies, we have used PPB staining and fluorescence microscopy to demonstrate that these signal voids correspond to the presence of iron-labeled cells in the brain [35, 36].

The size of the signal void created by iron-labeled cells is much larger than the actual cell size, due to what is known as the blooming effect. The blooming effect is a susceptibility artifact that occurs as a result of the iron oxide nanoparticle, which causes a local magnetic field inhomogeneity [34]. We have previously shown that we can detect single iron-labeled cells arrested in the mouse brain using cellular MRI [35, 39, 40]. Our image showed that the number of signal voids in the brain was similar for nude and NSG mice on day 0 and at endpoint. While we acknowledge that there is potential for immune cell uptake of the iron oxide nanoparticles that may be released by dead cancer cells, we believe that the majority of the signal voids that remain present are live cancer cells retaining iron. For example, in Parkins et al. [39] we investigated this mouse model using luciferase-positive 231BR cells and measured a strong correlation on day 0 between the number of signal voids detected in day 0 brain MR images and the brain signal measured in bioluminescence images, which is only detected from viable cells, providing evidence that signal voids represent live iron-labeled cells. In Hamilton et al. [41] we used fluorescence activated cell sorting (FACS) to successfully isolate live 231BR cells that were GFP-positive and labeled with red fluorescent (Flash Red) MPIO from the brains of mice. The GFP+ Flash Red+ cells were collected and expanded *in vitro*. The majority (~90%) of cells adhered to tissue culture plastic and successfully expanded, displaying the same cell morphology as the original cultured 231BR cells. This provide additional support for our claim that signal voids in MRI represent viable iron-positive cancer cells.

Our previous work has shown that the number of signal voids detected in MRI of the brain decreases over the course of the experiment, from day 0 to day 8, which is expected as the large majority of cancer cells that arrest in the brain do not survive and are cleared with time [39]. The signal voids which do persist in the brain over time are thought to represent iron-retaining, non-proliferative or dormant cancer cells. These cells have been shown to contribute to tumor recurrence [41]. It appears that both nude and NSG mice are capable of clearing dead cells from the brain.

3D MRI of the entire mouse brain at high resolution allowed us to view brain metastases in all 3 orientations and to digitally re-slice images to carefully interrogate image data. We counted and measured the volume of all MRI-detectable brain metastases. Brain metastases developed more quickly in NSG mice. At day 21 post cell injection NSG mice had more than 3 times the number of brain metastases. The NK cells and the remaining innate immune cells in nude mice likely contribute to the reduced tumor growth at this timepoint. This finding is in agreement with the results of other groups that show that more immune compromised mouse models are more permissive for tumor growth and metastasis [42–45]. Puchalapalli et al. [45] have previously reported an increase in the metastatic burden (in liver, lungs, brain and bones) in NSG compared to nude mice that were injected with the parental 231 breast cancer cell line in an intracardiac experimental metastasis model. In this study, metastases in each organ were enumerated from *ex vivo* fluorescence microscopy images. Our work demonstrates the advantages of using *in vivo* and longitudinal 3D MRI for the accounting of metastases in the whole brain.

NSG mice had to be euthanized at day 21/22. This was unexpected, as volumetric analyses of the tumor burden in the brain at this timepoint compared to the tumor burden of the nude mice at their endpoint of day 32 indicated that the NSG mice had lesser tumor burden in the brain. This suggested that there was a possibility of metastases to have occurred elsewhere in the body that could be contributing to a greater disease burden. Mouse body MRI confirmed tumors in the liver, lungs, and lymph nodes.

Nude mice were imaged at a third timepoint. On day 32 the number of brain metastases in nude mice was similar to the number of brain metastases in NSG mice on day 21. The mean volume of the brain metastases in the nude mice on day 32 was more than 3 times greater than those in NSG mice at day 21, having had more time to develop. Overall, the brain tumor burden at necessary endpoint was significantly greater in nude mice.

*Ex vivo* images of livers removed on the day of the intracardiac injection of iron labeled 231BR cells revealed numerous signal voids in all lobes of the liver in both strains of mice indicating that there was similar cancer cell arrest for both strains at this timepoint. We did not do a quantitative analysis of liver signal voids, however, in a previous study where we injected MPIO-labeled melanoma cells via the mesenteric vein, mouse livers were imaged *ex vivo* by MRI and the signal void area was shown to correlate with the number of cells injected [46]. *Ex vivo* images of livers removed at endpoint showed that numerous metastases formed in NSG mice while no liver metastases could be detected in nude mice.

The 231BR cell line was developed to selectively grow distant metastases in the brains of nude mice, and so the proliferation of metastases of this cell line in the liver demonstrates the loss of selectivity to the brain in the NSG mouse.

## Conclusion

In summary, high resolution cellular MRI allowed us to characterize the 231BR cell line in both the NSG and nude mouse models. We found marked differences in tumor incidence, volumes, and body tumor burden between strains. Our *in vivo* comprehensive analysis of cancer cell arrest, clearance, and tumor progression is important for understanding the metastatic cascade of a model of breast cancer brain metastasis that can be challenging to obtain with *in vitro* or *ex vivo* methods alone.

## List of Abbreviations

bSSFP: balanced steady-state free precession
BW: bandwidth
DMEM: Dulbecco’s modified Eagle’s medium
FA: flip angle
FACS: fluorescence activated cell sorting
FIESTA: Fast Imaging Employing Steady State Acquisition
GFP: green fluorescent protein
H&E: hematoxylin and eosin
MPIO: micron-sized iron oxide particles
MRI: magnetic resonance imaging
NSG: NOD/SCID/ILIIrg−/−
PBS: Perl’s Prussian Blue
PDX: patient-derived xenograft
RF: radiofrequency
TE: echo time
TR: repetition time
US: ultrasound
231BR: MDA-MB-231BR

## Declarations

### Ethics approval

Animals were cared for in accordance with the standards of the Canadian Council on Animal Care, and under an approved protocol of the Western University’s Council on Animal Care (2018-135).

### Consent to participate

Not applicable

### Consent for publication

Not applicable

### Availability of data and materials

The datasets used and/or analysed during the current study are available from the corresponding author on reasonable request.

### Code availability

Not applicable

### Author’s Contributions

P.J.F. designed the experiments. N.N.K, P.J.F, and A.M.H conducted the experiments. N.N.K. analyzed the data. N.N.K. and P.J.F. wrote the main manuscript text. All authors reviewed the manuscript.

## Acknowledgements

The authors would like to thank Ashley V. Makela for assisting with mouse injections for this study and the Canadian Institute for Health Research and the Breast Cancer Society of Canada for their funding.

